# Loss of the intellectual disability and autism gene *Cc2d1a* and its homolog *Cc2d1b* differentially affect spatial memory, anxiety, and hyperactivity

**DOI:** 10.1101/222638

**Authors:** Marta Zamarbide, Adam W. Oaks, Heather L. Pond, Julia S. Adelman, M. Chiara Manzini

**Affiliations:** Institute for Neuroscience and Department of Pharmacology and Physiology, The George Washington University School of Medicine and Health Sciences, Washington, DC, USA; Autism and Neurodevelopmental Disorders Institute, The George Washington University, Washington, DC, USA

**Keywords:** intellectual disability, learning, social function, anxiety, hyperactivity, rare diseases, mouse models

## Abstract

Hundreds of genes are mutated in non-syndromic intellectual disability (ID) and autism spectrum disorder (ASD), with each gene often involved in only a handful of cases. Such heterogeneity can be daunting, but rare recessive loss of function (LOF) mutations can be a good starting point to provide insight into the mechanisms of neurodevelopmental disease. Biallelic LOF mutations in the signaling scaffold *CC2D1A* cause a rare form of autosomal recessive ID, sometimes associated with ASD and seizures. In parallel, we recently reported that *Cc2d1a*-deficient mice present with cognitive and social deficits, hyperactivity and anxiety. In Drosophila loss of the only ortholog of *Cc2d1a, lgd*, is embryonic lethal, while in vertebrates *Cc2d1a* has a homolog *Cc2d1b* which appears to be compensating, indicating that *Cc2d1a* and *Cc2d1b* have redundant function in humans and mice. Here, we generate an allelic series of *Cc2d1a* and *Cc2d1b* loss of function to determine the relative role of these genes during behavioral development. We generated *Cc2d1b* knockout (KO), *Cc2d1a/1b* double heterozygous and double KO mice, then performed behavioral studies to analyze learning and memory, social interactions, anxiety, and hyperactivity. We found that *Cc2d1a* and *Cc2d1b* have partially overlapping roles. Overall, loss of *Cc2d1b* is less severe than loss of *Cc2d1a*, only leading to cognitive deficits, while *Cc2d1a/1b* double heterozygous animals are similar to Cc2d1a-deficient mice. These results will help us better understand the deficits in individuals with *CC2D1A* mutations, suggesting that recessive *CC2D1B* mutations and trans-heterozygous *CC2D1A* and *CC2D1B* mutations could also contribute to the genetics of ID.

## INTRODUCTION

Autosomal recessive loss of function (LOF) of the signaling scaffold <u>C</u>oiled-coil and <u>C2 D</u>omain containing <u>1A</u> (*CC2D1A*) causes a spectrum of neurodevelopmental conditions including fully penetrant intellectual disability (ID), and variably penetrant autism spectrum disorder (ASD), seizures, and aggressive behavior (Basel-Vanagaite et al., 2006; Manzini et al., 2014; Reuter et al., 2017). In *Drosophila*, where only one CC2D1 homolog, lethal giant discs *lgd*, is present, removal of *lgd* is lethal during the larval stage (Gallagher and Knoblich, 2006; Jaekel and Klein, 2006). Expression of either human CC2D1A or CC2D1B can rescue the phenotypes observed in *Drosophila* (Drusenheimer et al., 2015), suggesting that CC2D1A and CC2D1B act redundantly. Despite wide expression of CC2D1A and its binding to multiple proteins involved in the immune response (Chang et al., 2011; Chen et al., 2012), *CC2D1A* LOF in humans appears to only affect the brain, leading to a spectrum of behavioral deficits. While this indicates that CC2D1B is not fully able to compensate in the brain leading to the human presentation, it is unclear whether CC2D1B itself could have a role in neurodevelopmental disorders.

Studies on the genetic causes of ID and ASD, in particular, are identifying a large contribution of *de novo* and hypomorphic mutations to these diseases (Lim et al., 2013; Musante and Ropers, 2014; Sanders et al., 2012; Yu et al., 2013). Many of the mutated genes would have greater impact on development if completely lost, leading to multi-system disorders and/or brain malformations, while the heterozygous and hypomorphic mutations found in ASD/ID affect neurons more mildly, leading to a grossly normal brain, but with cognitive and social deficits (Yu et al., 2013). We wondered whether a similar mechanism is at play in patients with *CC2D1A* LOF mutations, where CC2D1B can only partially compensate. If this was the case, removal of both CC2D1 genes would be incompatible with embryogenesis, indicating that these proteins together have a critical developmental role. Nothing is known about the role of *CC2D1B* in brain development. By comparing how individual loss of each gene affects cognitive, social, and affective function we have studied the relative role of CC2D1A and CC2D1B in the brain and defined whether CC2D1B should also be considered as a candidate gene for ID.

Mice deficient for *Cc2d1a* develop normally *in utero*, but die soon after birth because of breathing and swallowing deficits (Al-Tawashi et al., 2012; Oaks et al., 2017; Zhao et al., 2011). By conditionally removing *Cc2d1a* in the forebrain we have previously shown that *Cc2d1a* LOF recapitulates features of ID and ASD in adult animals (Oaks et al., 2017). *Cc2d1a* conditional knockout (1a-cKO) mice show learning and memory deficits, social deficits, hyperactivity, anxiety, and repetitive behaviors (Oaks et al., 2017).

To define how CC2D1B compensates for loss of CC2D1A and contributes to these phenotypes, we generated a *Cc2d1b* knockout (1b-KO) line and developed an allelic series of *Cc2d1a* and *Cc2d1b* LOF, including *Cc2d1a/1b* double heterozygote (1a/1b-dHET) and double KO (1a/1b-KO) animals. Removal of both CC2D1 proteins causes early embryonic lethality, showing that CC2D1 function has an essential developmental role as in *Drosophila*. 1b-KO and 1a/1b-dHET animals are viable and fertile suggesting that *Cc2d1a* and *Cc2d1b* are not fully redundant, and that *Cc2d1a* has a critical role in respiration in the mouse.

When we tested the behavioral performance of 1b-KOs we found that *Cc2d1b* LOF caused only cognitive deficits, which are partially overlapping with those observed in *Cc2d1a* conditional LOF. Since direct comparison with a global *Cc2d1a* KO is not possible because of postnatal mortality, we also tested 1a/1b-dHETs which showed a combination of deficits with features of both 1b-KO and 1a-cKO animals, including delayed memory acquisition and retention, as well as increased anxiety and hyperactivity mostly in males. Our findings indicate that CC2D1 function is critical for embryonic development and that the CC2D1 proteins regulate multiple behaviors with some sex-specificity for males. Both CC2D1A and CC2D1B are involved in learning and memory, while CC2D1A alone appears to contribute to anxiety and hyperactivity.

## MATERIALS AND METHODS

### Animals

This study was carried out in accordance with the recommendations of the Institutional Animal Care and Use Committee of The George Washington University. A *Cc2d1b* null mouse line (1b-KO) was generated by the Knockout Mouse Project Repository (Project ID CDS 34981) at the University of California Davis, with the allele Cc2d1b^tm1a(KOMP)Wtsi^. *Cc2d1b* null mice carry an *engrailed 2* splice acceptor (En2SA) gene-trap allele with bicistronic expression of β-galactosidase as well as a neomycin resistance cassette, flanked by FRT (flippase recombinase target) recombination sites, in the genomic region between exons 2 and 3 of *Cc2d1b* (**Fig. 1A**). *Cc2d1a/1b* double heterozygous (1a/1b-dHET) mice were generated by crossing *Cc2d1a* heterozygotes (1a-HET) with *Cc2d1b* heterozygotes (1b-HET). 1a-HET mice were bred from a *Cc2d1a* null mouse line (KO) generated by the Knockout Mouse Project Repository (Project Design ID 49663) at the University of California as was previously described by Oaks et al (Oaks et al., 2017). All lines are maintained on a C57BL/6 background. For genotyping, polymerase chain reaction (PCR) amplifications were performed on 1*μ*L of proteinase K (New England Biolabs, Ipswich MA) digested tail DNA samples. PCR reactions (50*μ*L) consisted of GoTaq Flexi buffer (Promega, Madison WI), 100*μ*M dNTPs, 50*μ*M each of forward and reverse primers (sequence available upon request), 1mM MgCl_2_, and 1.25 U GoTaq Flexi DNA polymerase (Promega, Madison WI), and were run with optimized reaction profiles determined for each genotype. A 25*μ*L aliquot from each reaction was analyzed by gel electrophoresis on a 1.0% agarose gel for the presence of the desired band.

**Figure 1.**
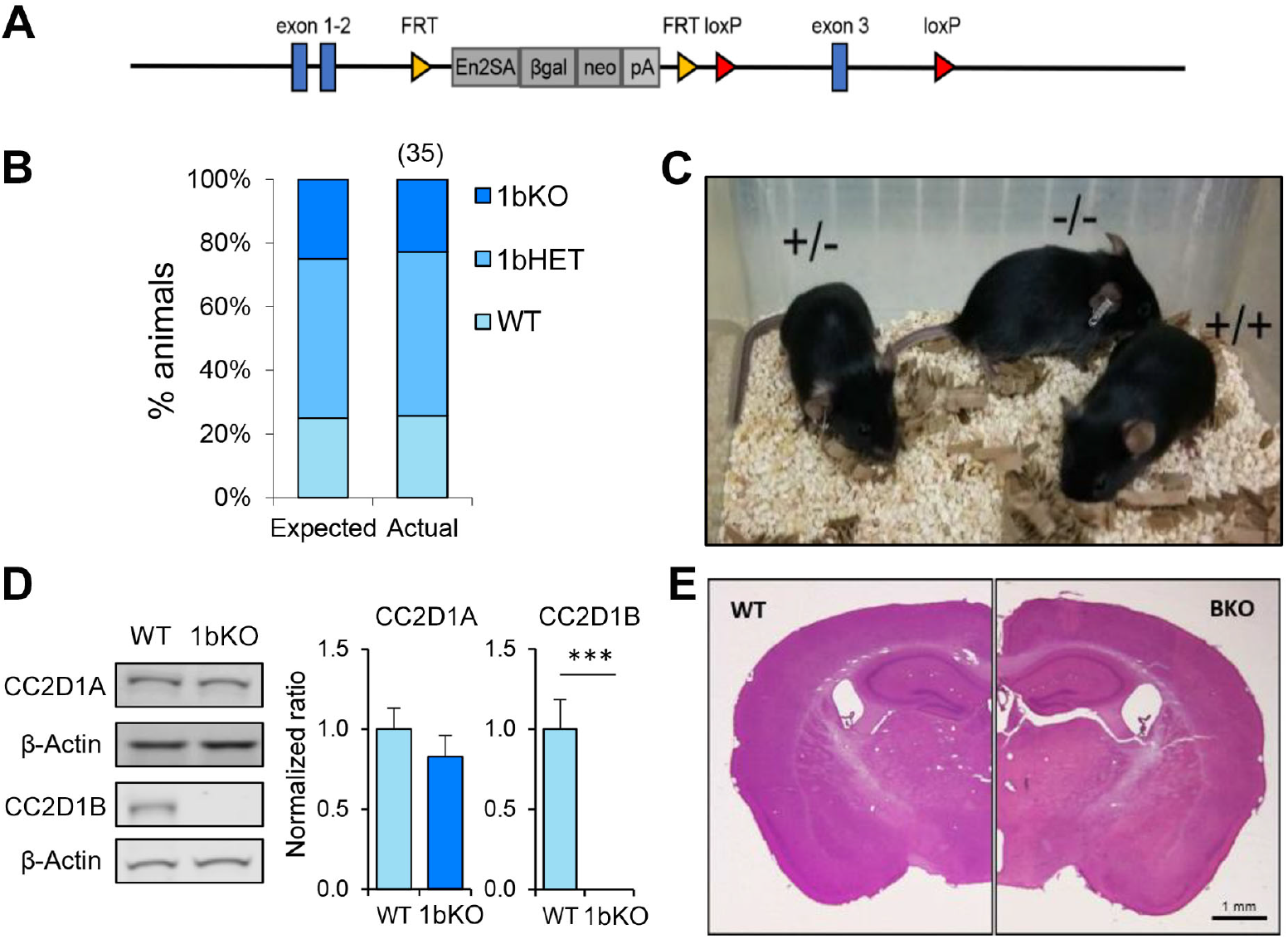
*Cc2d1b* KO mice are viable and fertile and present normal anatomical development of the brain. **A**. Gene trap containing engrailed 2 splice acceptor (En2SA) sequence followed by a β-Galactosidase cassette (βgal) and a neomycin resistance cassette (neo) is flanked by flippase recognition target (FRT) sites between exons 2 and 3. LoxP sites for Cre recombinase targeting flank exon 3. **B**. 1bKO mice are born in predicted Mendelian ratios (number of pups indicated above in parentheses; data from 5 litters) and **C**. are indistinguishable from WT and 1bHET mice. **D**. Immunoblot analysis of CC2D1A and CC2D1B expression in WT and 1bKO mice. Normal levels of CC2D1A and a complete absence of CC2D1B are shown in the 1bKO mice. (E) The size and organization of the adult 1bKO brain is indistinguishable from WT brain stained with hematoxylin and eosin. Scale bar: 1mm.

### Histological preparation and microscopy

To prepare tissue for histological analysis, deeply anesthetized mice were transcardially perfused with phosphate buffered saline (PBS) followed by 4% paraformaldehyde (PFA). Brains were removed and postfixed in PFA. Cryosections from adult mouse brains were prepared by mounting in Neg-50 (Thermo Scientific, Waltham MA) and cut at 40*μ*m on a Cryostar NX50 cryostat (Thermo Scientific, Waltham MA), then stained with Hematoxylin and Eosin (H&E, VWR International, Radnor PA) to visualize tissue architecture. Imaging of H&E stained sections was performed on a Leica M165 FC stereo microscope (Leica Microsystems, Buffalo Grove IL).

### Behavioral tests

A standardized battery of behavioral testing was applied to each cohort of animals, 1b-KO and 1a/1b-dHET male and female mice, at 3-4 months of age. As both 1b-KOs and 1a/1b-dHETs were generated from the same 1a/1b-dHET breeding the wild-type (WT) controls were littermates shared by both cohorts and all behavioral tests were performed at the same time for WT, 1b-KOs and 1a/1b-dHETs. Behavioral tests were performed in the Manzini lab behavioral suite in the George Washington University Animal Research Facility following a 60 min period of acclimatization. Initial characterization to analyze any neurological abnormalities included analysis of basic motor and somatosensory function was performed on a subset of the behavioral cohort as described by Rogers (Rogers et al., 1997): righting reflex, wire hang, gait analysis, tail pinch and visual reach. Cognitive and social function and other behaviors were tested in the open field test, novel object recognition test (Bevins and Besheer, 2006), Morris water maze (Vorhees and Williams, 2006), and 3-chamber social interaction test (Nadler et al., 2004). Behavioral analysis was performed via automated animal tracking using ANY-maze (Stoelting, Wood Dale IL).

#### Righting reflex

Coordination, motor strength and vestibular function were tested by placing each mouse on its back and timing its ability to return to an upright position.

#### Wire hang

Motor strength was tested by timing the latency to fall to a mouse cage containing bedding while the mouse was hanging from a wire cage-top not higher than 18 cm.

#### Gait analysis

Motor coordination and strength were assessed by painting the paws of each mouse with red non-toxic tempera paint and making them walk through a narrow tunnel over white paper. Abnormalities of paw placement and stride length were noted or indicated as normal.

#### Tail pinch

The ability of each mouse to respond to mild pain was tested by pinching the tip of the tail with fine, ethanol-cleaned forceps. Reactions were categorized as either response or no response.

#### Visual reach

Vision was tested by measuring the latency to the first attempt to reach for a nearby wire cage-top while the mouse was being held by the base of the tail at a height of 18 cm over an open cage.

#### Open field test

The open field test was performed in an unfamiliar 50×50 cm plastic box (Stoelting, Wood Dale IL). Animals were placed in the center of the arena and ambulatory activity was monitored by digital video for 15 min. The arena was divided into two areas, an outer zone and a center zone (25×25 cm; 25% of total area). Total distance traveled and time spent in each area were measured.

#### Novel object recognition test

The novel object recognition test (Bevins and Besheer, 2006; Oaks et al., 2017) was performed in the same apparatus described for the open field test. The test consisted of three different phases: habituation, training and test. The habituation phase lasted for 30 min while the animals were exposed to the box and then returned to the home cage while the box was cleaned. During the training phase, the animal was placed in the same box with two identical objects located in opposite corners, at a distance of five cm from the walls. To assess short-term memory, the animal was returned to the home cage during an interval of 15 min. During the test, a familiar object, identical to those used in the habituation phase, was placed in one corner, while in the opposite corner an unfamiliar object was placed. Exploration activity was monitored for 10 min at each phase, with exploration defined as time spent actively observing or touching the object from within a radius of five cm. Cumulative time spent with each object was measured by video analysis using ANY-maze to determine the location of the animal’s nose relative to the objects in the enclosure. Preference for the novel object was defined as the ratio of the time spent with the novel object to the time spent with the familiar object. Animals that did not interact with the object and stopped in a corner of the cage were removed from the analysis.

#### Morris water maze

The Morris water maze (Oaks et al., 2017; Vorhees and Williams, 2006) apparatus was a 120x120cm round metal tub (Stoelting, Wood Dale IL) where distinct visual cues were placed at the cardinal points. White non-toxic paint was added to the water to make the surface opaque for the hidden trials and it was maintained at 24 °C. Each trial consisted of four independent drops, one at each cardinal point around the tub, with the mouse facing the wall of the tub. Each drop lasted 60s, or until the mouse found the platform, whichever occurred first. Each animal completed two trials (four drops each) with a visible platform, five trials with a platform hidden under the water surface, and two reversal trials where the location of the hidden platform was changed. The sequence of nine trials was performed over nine days, with one trial per day. A 60 s probe trial was also performed the day after the hidden platform series was completed, by removing the platform from the water before proceeding to the reversal phase on the following day.

#### Three-chamber social interaction test

The social interaction test (Kaidanovich-Beilin et al., 2011; Nadler et al., 2004) was performed in a clear rectangular acrylic box (60x40 cm) divided into three chambers (40x20 cm) with small openings (10x5 cm) in the adjoining walls (Everything Plastic, Philadelphia PA). The test consisted of two phases, the habituation and the sociability phase. During the habituation phase, empty inverted wire cups (10 cm diameter) were placed in the center of the chambers at the ends. Each mouse was placed in the center chamber of the apparatus and allowed to explore the different chambers for 5 min. During the second phase, an unfamiliar mouse of the same sex as the tested mouse was placed under the wire cup in one of the side chambers. The experimental mice were allowed to explore for 10 min during the sociability phase. Total time spent in the Object (containing empty cup) and Mouse (with unfamiliar mouse under the cup) chambers was used to determine the social preference of each mouse tested, while the time sniffing within a 2-cm radius of the mouse-containing cup were recorded as measures of social approach and social interaction.

## RESULTS

### CC2D1A and CC2D1B have partially redundant function in development

Loss of *CC2D1A* in humans causes a variable spectrum of ID, ASD and seizures and removal of *Cc2d1a* in the murine forebrain leads to several cognitive, social, and affective behavioral phenotypes (Manzini et al., 2014; Oaks et al., 2017). As no human mutations in *CC2D1B* have been identified to date, we asked whether loss of *Cc2d1b* in the mouse would lead to similar phenotypes as loss of *Cc2d1a*. A *Cc2d1b*-deficient line (1b-KO) had been generated from the Knockout Mouse Project (KOMP) as a gene-trap allele inserted in intron 2 of *Cc2d1b* (**Fig. 1A**). We obtained heterozygous animals and bred them to homozygosity, finding that 1b-KO mice are born in Mendelian ratios (**Fig. 1B**). Differently from *Cc2d1a* KO (1a-KO) pups, which die shortly after birth (Al-Tawashi et al., 2012; Drusenheimer et al., 2015; Oaks et al., 2017; Zhao et al., 2011), 1b-KO mice are viable, fertile, and are indistinguishable from WT littermates (**Fig. 1C**). Basic behavioral functions were tested in adult WT and 1b-KO males and females: coordination (righting reflex), strength (wire hang), locomotion (stride and gait), pain sensitivity (tail pinch), and vision (visual reflex). No differences were observed in basic sensory and motor function (**Table 1**). We confirmed via Western blot analysis of cortical protein lysates that CC2D1B was completely absent in these animals and that CC2D1A was expressed at normal levels (**Fig. 1D**). Cryosections generated from the adult brain of 1b-KO animals and stained using hematoxylin and eosin (H&E) showed no differences in brain size and organization from WT littermates (**Fig. 1E**). In summary, loss of *Cc2d1b* does not affect respiratory function and deglutition in the infant as observed in 1a-KOs, and 1b-KO adult mice are indistinguishable from WT littermates.

**Table 1.** Analysis of basic motor and sensory function in 1b-KO and 1a/1b-dHET mice

| Genotype | Sex | Weight (g) | Righting reflex (s) | Wire hang (s) | Stride and gait | Tail pinch<sup>a</sup> | Visual reflex<sup>b</sup> |
| --- | --- | --- | --- | --- | --- | --- | --- |
| WT | M | 28.39 ± 0.94 | <1 s | 56.5 ± 2.35 | Normal | 6/9 | 9/9 |
| 1b-KO | M | 27.49 ± 0.67 | <1 s | 52.21 ± 5.05 | Normal | 7/10 | 10/10 |
| 1a/1b-dHET | M | 28.32 ± 0.85 | <1 s | 53.61 ± 3.24 | Normal | 6/8 | 8/8 |
| WT | F | 21.06 ± 0.53 | <1 s | 58.7 ± 1.09 | Normal | 7/10 | 10/10 |
| 1b-KO | F | 21.95 ± 0.83 | <1 s | 55.87 ± 3.71 | Normal | 6/8 | 8/8 |
| 1a/1b-dHET | F | 21.81 ± 0.88 | <1 s | 58.2 ± 1.59 | Normal | 7/10 | 10/10 |

<sup>a</sup> Animals responding to tail pinch. <sup>b</sup> Animals responding to visual stimulus.

CC2D1A and CC2D1B contain very similar protein domains and are thought to have redundant functions in endocytic traffic and gene transcription (Drusenheimer et al., 2015; Hadjighassem et al., 2009; Usami et al., 2012). Because CC2D1B loss of function did not result in postnatal lethality, we wondered whether the two proteins would only be partially redundant. To test this hypothesis, we crossed 1b-KOs and 1a-KOs to generate *Cc2d1a/Cc2d1b* double heterozygous (1a/1b-dHET) and double KO mice (1a/1b-KO). As 1aKO pups die soon after birth (Al-Tawashi et al., 2012; Oaks et al., 2017; Zhao et al., 2011), we did not expect 1a/1b-KO animals to survive and we genotyped litters at postnatal day (P)0, collecting tissue from both live and dead pups. However, while dead 1a-KO and 1a-KO/1b-HET were found in the expected ratios, 1a/1b-dKO pups were never retrieved (**Fig. 2A**), suggesting that double knockouts may die earlier during embryonic development. Examination of prenatal litters only identified 1a/1b-dKO tissue mid-gestation at E11.5, but the embryo was almost entirely absent, leaving only a hypomorphic and largely empty yolk sac (**Fig. 2B**). These results indicate that removal of both CC2D1 proteins leads to early embryonic lethality.

**Figure 2.**
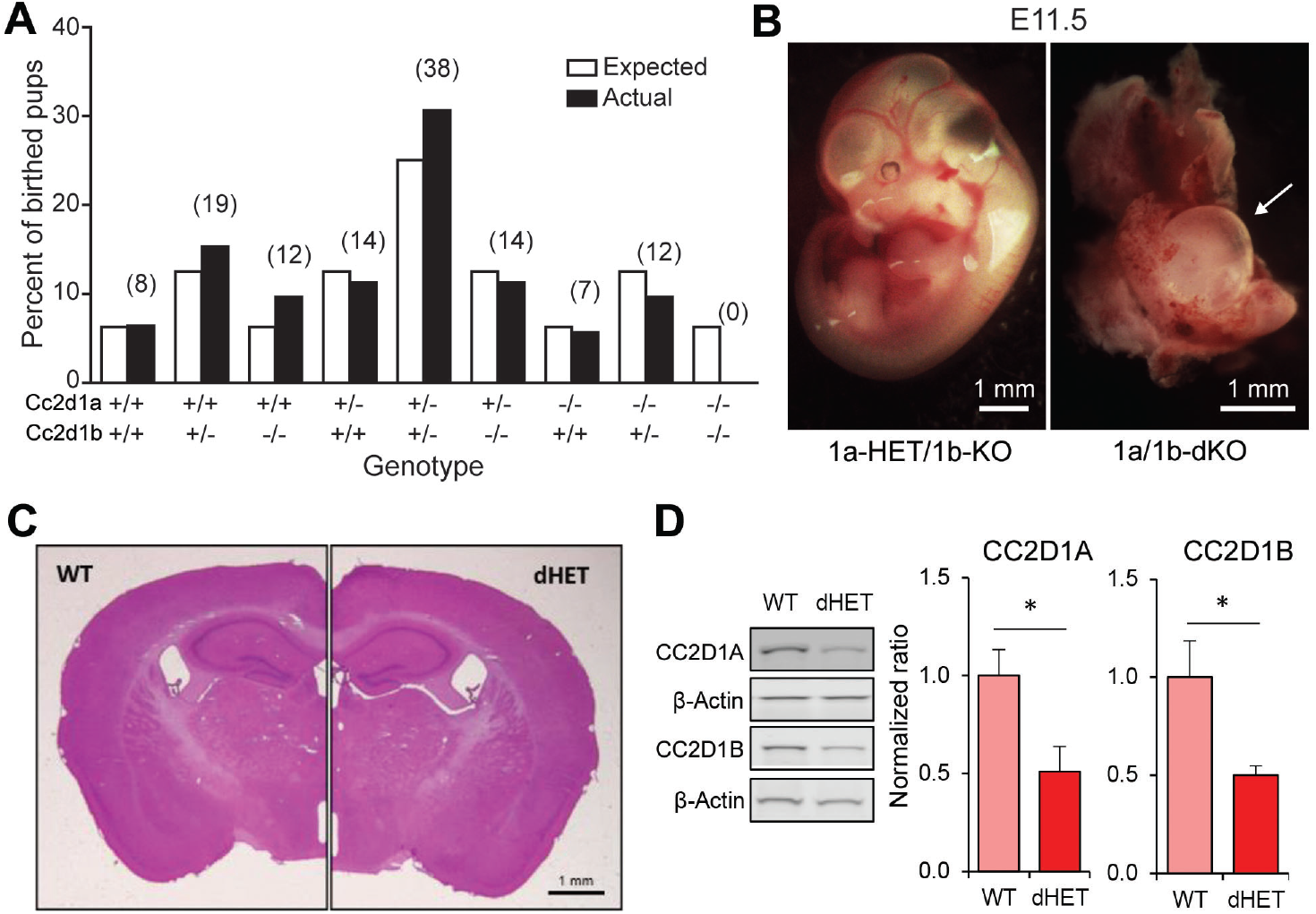
*Cc2d1a/Cc2d1b* double LOF is embryonic lethal, while double heterozygotes are viable. **A**. Genotypes at postnatal day 0 (P0) of 124 pups resulting from 22 double heterozygous (1a/1b-dHET) crosses (number of pups for each genotype indicated above in parentheses). The 1a/1b double knockout (A-/-B-/-) pups were never found at P0. **B**. Representative images of normal embryonic day 11.5 (E11.5) embryo with a single intact *Cc2d1* allele (left panel) and a double KO embryo (right panel; arrow indicates empty yolk sac). Scale bars: 1mm **C**. The size and organization of the adult 1a/1b-dHET brain is indistinguishable from wild type mice stained with hematoxylin and eosin. Scale bar: 1mm **D**. Immunoblot analysis of CC2D1A and CC2D1B expression in wild-type and 1a/1b-dHET mice. A half dose of each CC2D1 protein was found. Results expressed as mean ± SEM. *p<0.05 (two tailed t-test).

1a/1b-dHETs were viable and fertile and indistinguishable from WT littermates with normal gross brain anatomy (**Fig. 2C**), and normal basic motor and sensory function (**Table 1**). We tested the expression levels of CC2D1A and CC2D1B in 1a/1b-dHET mice and found that as expected only a half dose of each CC2D1 protein was present (**Fig. 2D**). Thus, combined CC2D1 function is necessary for embryonic morphogenesis, but 1b-KO or 1a/1b-dHET animals develop normally, indicating that CC2D1A and CC2D1B have similar functions as it pertains to gross anatomical development and survival.

### Both CC2D1A and CC2D1B are important for cognitive function

We have previously found that loss of *Cc2d1a* leads to a constellation of behavioral deficits: cognitive and social impairment, anxiety, hyperactivity and repetitive behaviors (Oaks et al., 2017). We generated a cohort of 1b-KO and 1a/1b-HET male and female mice for behavioral analysis by crossing 1a/1b-HETs, so that we could compare behavioral performance in both lines at the same time. In the short-term memory version of the Novel Object Recognition Test (NORT) (Bevins and Besheer, 2006) mice are placed in an arena with two identical objects that they are free to explore. After being removed back to their cages for 15 minutes, they are put in the arena where one of the now known objects has been substituted for a novel object (**Fig. 3A**). In this test, WT male and female mice spend roughly four times longer exploring the novel object, while 1b-KOs and 1a/1b-dHETs show no difference (**Fig. 3B–C**) (Males: WT, T2/T1=1.21±0.32, T4/T3=3.90±0.75, n=10, p=0.004 **; 1b-KO, T2/T1=1.05±0.24, T4/T3=1.60±0.46, n=11, p=0.309; 1a/1b-dHET, T2/T1=1.08±0.23, T4/T3=1.62±0.46, n=12, p=0.307. Females: WT, T2/T1=1.20±0.25, T4/T3=4.39±1.40, n=10, p=0.038 *; 1b-KO, T2/T1=0.84±0.16, T4/T3=0.93±0.24, n=10, p=0.757; 1a/1b-dHET, T2/T1=1.34±0.48, T4/T3=1.46±0.28, n=10, p=0.824). This deficit was not due to reduced interest in the objects, as animals spent similar amounts of time in exploratory behaviors, with 1a/1b-dHET males showing significantly more exploration (**Fig. 3D**. T1+T2 - Males: WT, t=26.97±5.75s, n=10; 1b-KO, t=23.17±3.65s, n=11, p=0.999; 1a/1b-dHET, t=65.27±15.93s, n=12, p=0.167; Females: WT, t=40.56±5.19s, n=10; 1b-KO, t=71.67±17.47s, n=10, p=0.423; 1a/1b-dHET, t=54.30±10.56s, n=10, p=0.960. **Fig. 3E**. T3+T4 - Males: WT, t=21.93±5.54s; 1b-KO, t=17.91±3.57s, p=0.999; 1a/1b-dHET, t=50.83±16.0s, p=0.640. Females: WT, t=15.39±2.12s; 1b-KO, t=68.38±26.04s, p=0.090; 1a/1b-dHET, t=31.86±10.61s, p=0.959. **Fig. 3F**. SUM T1,2,3,4 - Males: WT, t=48.90±9.35s; 1b-KO, t=41.08±6.20s, p=0.942; 1a/1b-dHET, t=116.1±28.24s, p=0.033 *; Females: WT, t=55.95±6.62s, 1b-KO; t=140.1±42.64s, p=0.073; 1a/1b-dHET, t=86.16±20.52s, p=0.660).

**Figure 3.**
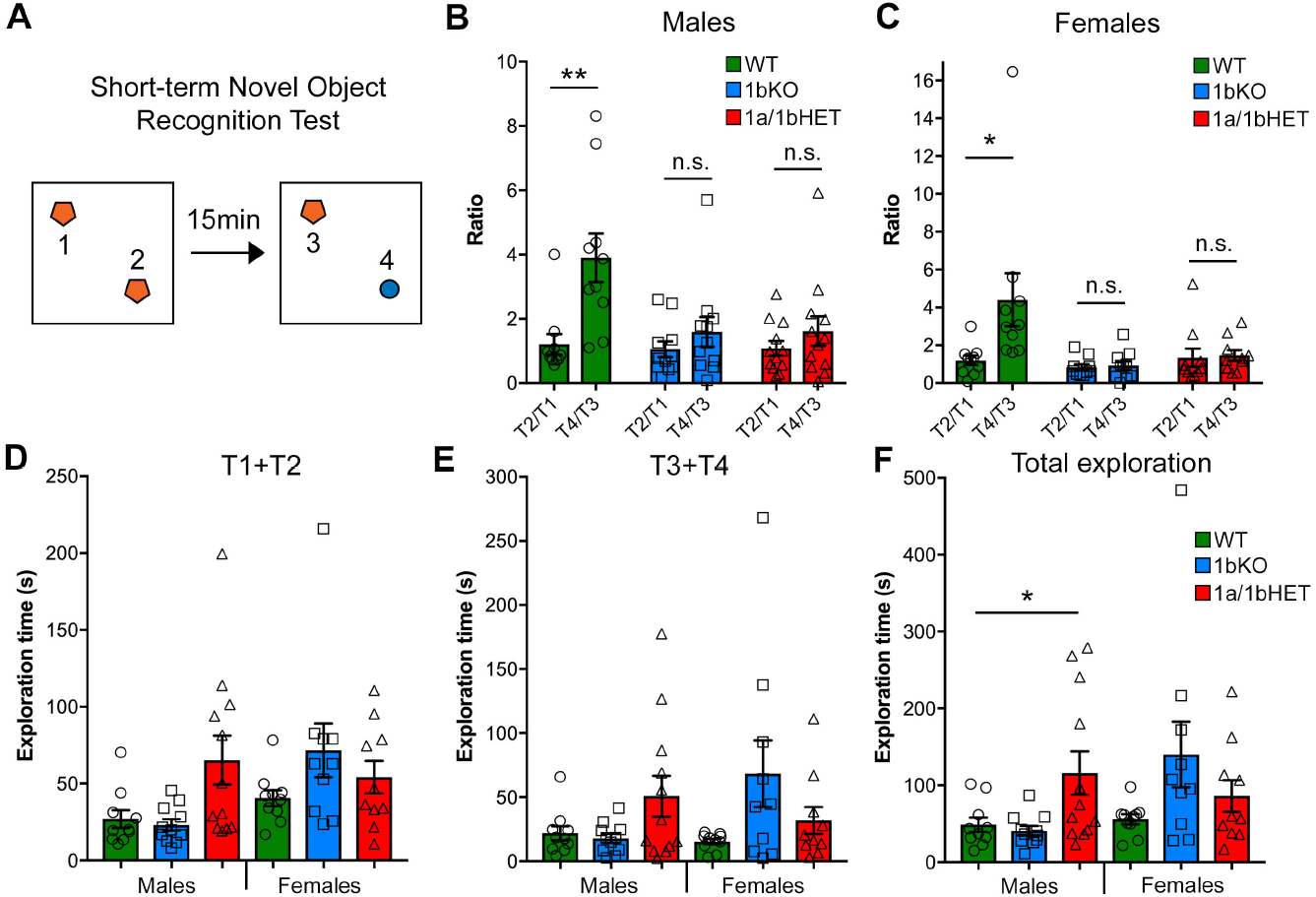
CC2D1A and CC2D1B are both involved in object memory. **A**. Schematic design of short-term novel object recognition test (NORT), with a novel object replacing a familiar object after a 15 min interval. **B-C**. In contrast to WT, 1bKO and 1a/1bHET male (**B**.) and female (**C**.) mice showed no preference for the novel object relative to a familiar object. Results expressed as mean ± SEM, **p<0.01 (two-tailed t-test with equal variance). **D-F**. Exploration time divided by initial exploration of training objects 1 and 2 (**D**.), test objects 3 and 4 (**E**.) and total exploration across the two phases of the NORT (**F**.). 1a-1bHET males show a trend towards increased exploration in each test phase which reaches significance when both phases are combined. Results expressed as mean ± SEM, One-way ANOVA with Dunnett’s multiple comparison test, *p<0.05, **p<0.01

To further assess cognitive function, the 1b-KO mice were tested using the Morris Water Maze (MWM) paradigm which probes spatial memory acquisition, retention and flexibility, by testing the ability of a mouse to learn, remember, and relearn the location of a platform hidden under opaque water (Morris, 1984). After the mice are trained using a visible platform to escape from the water, the platform is hidden under the surface in a different location and the animals undergo training on five consecutive days to learn the location of the platform. On the following day, memory retention is tested by removing the platform and measuring the amount of time the mouse spends in the area where the platform was previously located (probe trial). Finally, the position of the platform is changed and the animal must display flexibility by learning a new location (reversal). 1a-cKO animals show a delay in initial acquisition of the location of the hidden platform (HP), but after they learn, they can retain the memory in the probe trial, and learn a new location in the reversal (Oaks et al., 2017). Both 1b-KO and 1a/1b-dHET males and females presented deficits in this test (**Fig. 4**). 1b-KO males and females and 1a/1b-dHET males were delayed in the hidden platform acquisition showing significant differences in day 2 or 3 of the test (HP2 and HP3 in **Fig. 4B and F**) (Males HP3: WT, t=6.82±0.69s, n=11; 1b-KO, t=10.97±1.85s, n=10, p=0.042 *; 1a/1b-dHET, t=11.99±1.28s, n=13, p=0.0027 **. Females HP2: WT, t=12.30±1.32s, n=13; 1b-KO, t=19.62±1.74s, n=10, p=0.0025 **; 1a/1b-dHET, t=14.66±1.64s, n=11, p=0.247). 1a/1bHET males and females were also affected in the probe trial where they spent less time in the platform quadrant during the first 15sec of the 60sec trial (**Fig. 4D and H**) (Probe 15s - Males: WT, t=9.51±0.83s, n=11; 1b-KO, t=6.13±0.50s, n=10, p=0.0029 **; 1a/1b-dHET, t=5.48±0.80s, n=13, p=0.0021 **. Females: WT, t=7.18±0.80s, n=13; 1b-KO, t=5.77±0.65s, n=10, p=0.203; 1a/1b-dHET, t=4.36±0.82s, n=11, p=0.022 *). Finally, 1b-KO males, but not females, were affected throughout the 60sec probe trial and spent less time exploring the correct quadrant in the probe trial testing memory retention (**Fig. 4D**) (Probe 60s - Males: WT, t=25.40±1.78s, n=11; 1b-KO, t=19.58±1.30s, n=10 p=0.018 *; 1a/1b-dHET, t=22.74±2.63s, n=13, p=0.428. Females: WT, t=21.19±1.85s, n=13; 1b-KO, t=20.57±1.54s, n=10, p=0.809; 1a/1b-dHET, t=18.43±2.62s, n=11, p=0.389). Animals heterozygous for loss of *Cc2d1a* or *Cc2d1b* alone showed normal behavioral performance (**Suppl. Table 1** and **Fig. 1–2**). In summary, loss of CC2D1B lead to cognitive deficits in both memory acquisition and retention. In general, males appear more severely affected than females in both 1bKO and 1a/1bHET lines, suggesting that CC2D1A and CC2D1B have overlapping roles in cognitive function.

**Figure 4.**
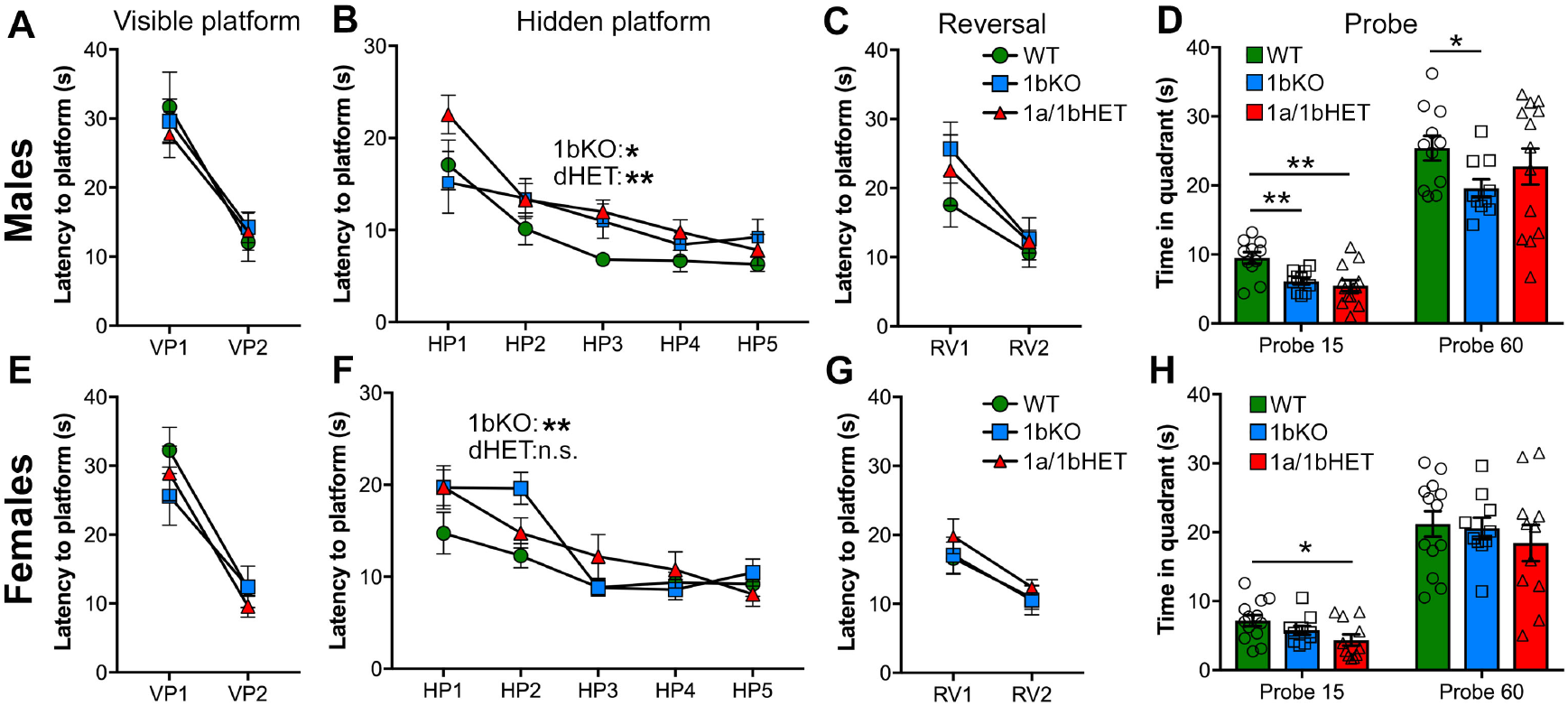
CC2D1B is involved in spatial memory formation and retention with mild male-specificity. Hippocampus-dependent spatial memory was assessed in 1bKO and 1a/1bdHET mice via the Morris Water Maze test. Spatial learning was measured as latency to escape in three different stages, visible platform (VP), hidden platform (HP), or the reversal (RV) of the hidden platform position. No deficits were shown by males (**A**.) or females (**E**.) of any genotype in identifying the platform in the VP trial. **B**. Both 1bKO and 1a/1bHET males showed a delay in learning the location of the hidden platform, and a similar deficit was present in 1bKO females (**F**.). **C**. and **G**. No differences were found in the RV during the test. **D**. and **H**. Spatial memory retention was measured between the HP and RV trials by the time spent swimming in the quadrant where the platform was previously located. Significant spatial memory impairment was found in the 1bKO male mice compared to WT both during the first 15sec and at the end of the trial after 60sec, while female 1bKO mice showed no deficit. 1a/1bHET males and females spent less time looking for the platform during the first 15sec, but subsequently recovered. Two-way ANOVA with repeated measures was used for analysis of the HP phase. Multiple t-tests with equal variance were uses for individual timepoints and probe analysis *p<0.05, **p<0.01

### Only CC2D1A is involved in anxiety and hyperactivity

1A-cKO animals showed increased mobility and reduced entry into the center of the open field arena, indicating hyperactivity and anxiety (Oaks et al., 2017). In addition, removal of *Cc2d1a* in the forebrain also leads to ulcerative dermatitis due to obsessive grooming and social interaction deficits (Oaks et al., 2017). 1b-KO males and females performed similarly to WT littermates in the open field test and showed no signs of hyperactivity or anxiety (**Fig. 5**) (Distance - Males: WT, d=25.16±2.29m, n=11; 1b-KO, d=29.63±1.96m, n=11, p=0.498; Females: WT, d=34.65±1.36m, n=13; 1b-KO, d=42.37±3.28m, n=11, p=0.097. Time in center - Males: WT, t=78.13±5.23s, n=11; 1b-KO, t=83.17±14.26s, n=11, p=0.988; Females: WT, t=77.45±11.78s, n=10; 1b-KO, t=87.75±17.65s, n=10, p=0.969). Interestingly, 1a/1b-dHETs showed increased locomotion and avoidance of open spaces, as previously observed for the 1a-cKOs, but only in males similarly to the exploration in the NORT where increased exploratory behavior was only observed in 1a/1b-dHET males (**Fig. 5A-B**) (Distance - Males: WT, d=25.16±2.29m, n=11; 1a/1b-dHET, d=35.85±2.94m, n=13, p=0.0076 **; Females: WT, d=34.65±1.36m, n=13; 1a/1b-dHET, d=35.35±2.51m, n=11, p=0.999. Time in center - Males: WT, t=78.13±5.23s, n=11; 1a/1b-dHET, t=41.32±3.71s, n=13, p=0.0198 *; Females: WT, t=77.45±11.78s, n=10; 1b-KO, t=121.90±15.19s, n=11, p=0.1225). No ulcerative dermatitis or obsessive grooming was observed in any of these mouse lines.

**Figure 5.**
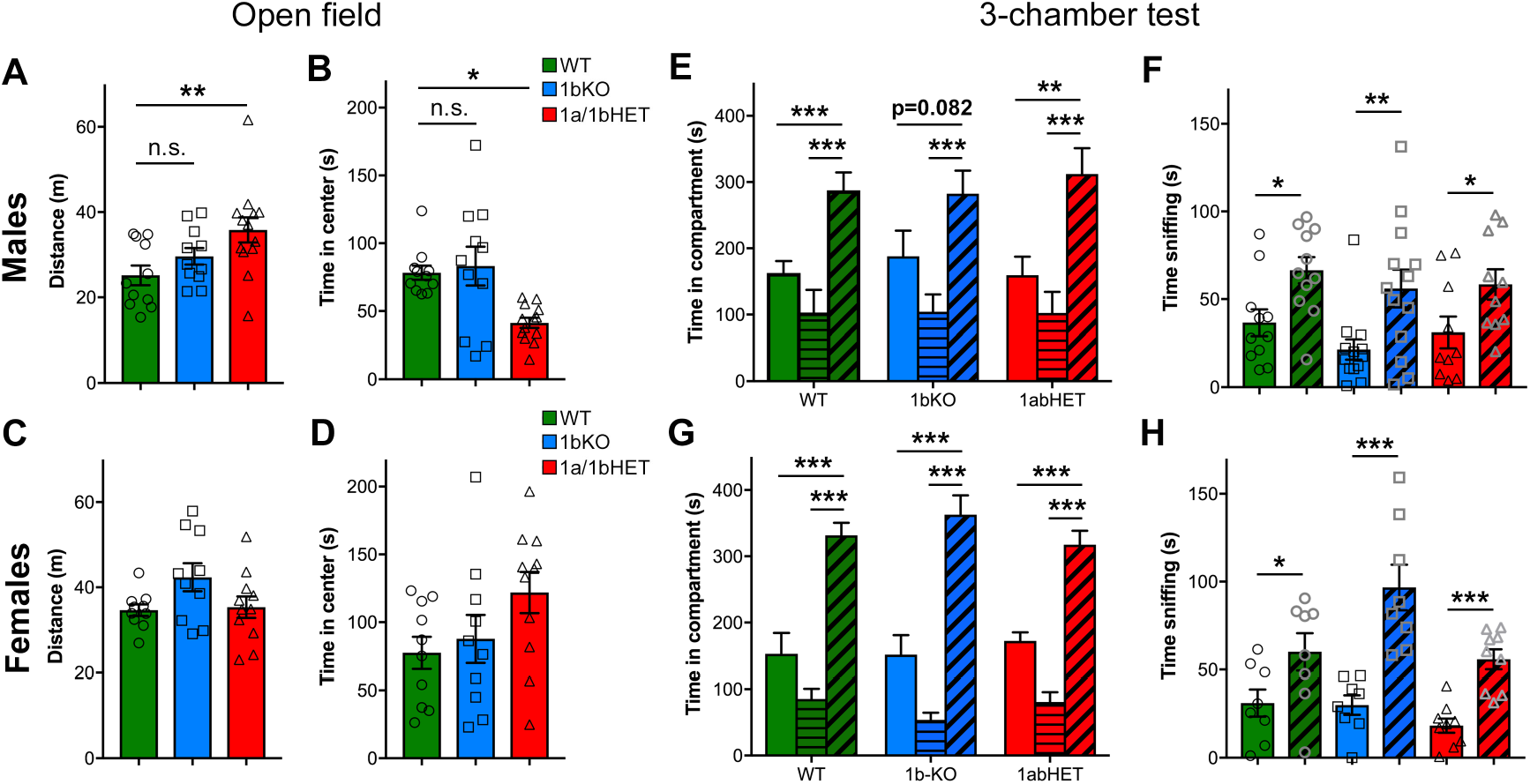
CC2D1A contributes to anxiety and hyperactivity. **A-D**. Exploratory and general locomotor activity in a novel environment was assessed on the open field test. Total path length (**A**.) and time spent in the center zone (**B**.) are only affected in 1a/1bHET male mice, while females show no difference (**C-D**.). Results expressed as mean ± SEM, One-way ANOVA with Dunnett’s multiple comparison test, *p<0.05, **p<0.01 **E-H**. Social interaction behavior was assessed by the three-chamber test, presented as time spent in each chamber (**E.,G**.) and time spent sniffing the novel mouse vs. the empty cup (**F.,H**.). 1bKO male mice showed significantly increased sniffing of the novel mouse vs. the cup (**F**.), but did not show a significant increase in time in the compartment indicating that they may display less interest for the mouse (**E**.). 1a/1bHET males and females and 1bKO females showed no difference from WT littermates (**E-H**.). Results expressed as mean ± SEM. Two-tailed t-test with equal variance *p<0.05, **p<0.01, ***p<0.001

Finally, all mice were tested in the social approach version of the three-chambered test. In this test, the mouse is placed in an apparatus with three communicating chambers. In the left chamber, there is a novel mouse of the same sex under a wire cup, while in the right chamber there is an empty wire cup. Mice spend more time exploring and sniffing the stranger mouse than the object and this is considered a social action (Kaidanovich-Beilin et al., 2011; Nadler et al., 2004). The 1a-cKO showed no preference for the conspecific both as in the time spent around the mouse enclosure and the time spent sniffing the stranger mouse (Oaks et al., 2017). 1a/1b-dHET males and females and 1b-KO females behaved like WT mice in this test (**Fig. 5E-H**). 1b-KO males were moderately affected showing non-significant difference between the empty cup and the stranger (**Fig. 5E**) [Males: WT, time with mouse (tm)=287.65±26.81s, time with object (to)=162.74±18.15s, n=11, p=0.00098 ***; 1b-KO, tm=282.37±34.83s, to=187.98±28.63s, n=13, p=0.082; 1a/1b-dHET, tm=312.05±39.03s, to=159.39±28.11s, n=10, p=0.0052 **. Females: WT, tm=331.50±19.14s, to=152.70±31.59s, n=8, p=0.00026 ***; 1b-KO, tm=362.93±29.06s, to=151.53±29.39s, n=8, p=0.00016 ***; 1a/1b-dHET, tm=317.62±20.89s, to=172.47±12.91s, n=9, p=0.00002 ***]. The deficit in 1b-KO males was primarily due to a subset of animals showing preference for the object (**Suppl. Fig. 3**). All genotypes showed significantly increased time spent sniffing the stranger mouse, indicating that once in the chamber the 1b-KO animals interact with the other animal (**Fig. 5F and H**) [Males: WT, time sniffing mouse (tsm)=66.47±7.44s, time sniffing object (tso)=36.61±7.51s, n=11, p=0.0105 *; 1b-KO, tsm=56.04±10.78s, tso=21.47±5.78s, n=13, p=0.009 **; 1a/1b-dHET, tsm=58.40±8.65s, tso=31.11±8.95s, n=10, p=0.042 *. Females: WT, tsm=60.11±10.60s, tso=31.15±7.71s, n=8, p=0.044 *; 1b-KO, tsm=96.68±13.00s, tso=29.93±5.55s, n=8, p=0.00033 ***; 1a/1b-dHET, tsm=55.80±5.66s, to=18.26±4.02s, n=9, p=0.00005 ***].

In conclusion, 1b-KO and 1a/1b-dHET animals show only partially overlapping behavioral profiles in anxiety, hyperactivity, and sociability. 1b-KO mice of either sex do not appear anxious or hyperactive and only males show a mild sociability deficit in the three-chamber test. 1a/1b-dHET males are more similar to 1a-cKO mice with increased locomotion and decreased time in the center of the open field. These results show that CC2D1A and CC2D1B only have partially redundant roles in cognitive and social function. Each of the *Cc2d1* genes contributes to aspects of learning and memory and sociability, but *Cc2d1a* appears to be more critical for hyperactivity and anxiety. Interestingly, both lines display sexually dimorphic phenotypes with males being mildly more affected than females.

## Discussion

Cognitive development is controlled by a multitude of mechanisms regulating synaptic transmission and neuronal function. Hundreds of genes have been found mutated in patients with ID and ASD and the generation of mouse models has deepened our understanding of how each gene contributes to disease and behavior (Ey et al., 2011; Kazdoba et al., 2015; Nestler and Hyman, 2010). Mutations in the gene encoding CC2D1A cause a rare form of ID and ASD in humans, and this protein is emerging as a critical regulator of intracellular signaling with roles in cognitive function (Basel-Vanagaite et al., 2006; Manzini et al., 2014), immunity (Chang et al., 2011; Zhao et al., 2010) and cancer (Yamada et al., 2015). Removal of the only *CC2D1* homolog in Drosophila, *lgd*, causes early lethality and severe deficits in morphogenesis, and both human proteins can rescue *lgd* LOF phenotypes, suggesting that the vertebrate CC2D1 proteins have redundant functions (Drusenheimer et al., 2015). In fact, deficits in *lgd* mutant flies are more severe than in 1a-KO and 1b-KO mice (Drusenheimer et al., 2015). We hypothesized that the neuropsychiatric phenotypes observed in humans carrying *CC2D1A* LOF mutations are likely due to the inability of CC2D1B to fully substitute for CC2D1A.

Initial evidence to support our hypothesis was provided by the fact that 1a-KO mice are anatomically normal but die soon after birth due to breathing and swallowing deficits (Al-Tawashi et al., 2012; Chen et al., 2012; Oaks et al., 2017; Zhao et al., 2011), while 1b-KOs are viable and fertile (Drusenheimer et al., 2015). No respiratory deficits have been reported in humans with *CC2D1A* mutations and these findings indicated that *Cc2d1a* has an essential role in breathing regulation in the brain stem in the mouse where CC2D1B cannot complement CC2D1A function. We do not know whether this difference between mice and humans is due to the timing of birth which is at an earlier stage of neural development in mice, or to differences in CC2D1A and CC2D1B expression in the brain stem in the two species.

The current study provides further evidence that *Cc2d1a* LOF is more severe than *Cc2d1b* LOF through behavioral studies. Forebrain-specific Cc2d1a-deficient mice 1a-cKO display an array of cognitive and social deficits, in addition to anxiety and hyperactivity (Oaks et al., 2017). 1b-KO mice only display cognitive deficits, with object recognition impairment in the NORT and reduced memory acquisition and retention in the MWM test, but no other phenotypes. Interestingly, the MWM test results reveal different roles for the CC2D1 proteins in spatial learning and memory. 1a-cKO animals showed delayed learning, but no deficit in remembering the location of the platform once it was learned (Oaks et al., 2017), while 1b-KO mice also displayed reduced memory retention in the probe especially in males. Parallel studies in the 1a/1b-dHET line confirm this difference observing deficits in both spatial memory acquisition and retention. In comparing cognitive performance in 1b-KOs with 1a/1b-dHET and previously published 1a-cKOs, all lines were equally deficient in the NORT, indicating that object recognition circuits in the cortex and hippocampus are affected (Antunes and Biala, 2012).

*Cc2d1b* also differs from *Cc2d1a*, as it appears to have no role in social behavior, hyperactivity and anxiety. Results from the 1a/1b-dHETs suggest that partial loss of *Cc2d1a* in combination with a half dosage of *Cc2d1b* is sufficient to generate hyperactivity and anxiety. Interestingly, only complete loss of *Cc2d1a* leads to social deficits. Taken together, our results indicate that *Cc2d1a* and *Cc2d1b* have roles in behavioral function that are only partially redundant. Behavior is regulated by a multitude of molecular and cellular mechanisms, but it is interesting to note how each of these two homologous proteins may contribute to specific sets of behaviors. These effects could be due to their role in controlling a variety of intracellular signaling processes and thereby affecting multiple cellular functions.

CC2D1A and CC2D1B were reported to regulate endocytosis and gene transcription (Drusenheimer et al., 2015; Hadjighassem et al., 2011; Martinelli et al., 2012; Usami et al., 2012), but CC2D1A has been the most studied to date. Many of the pathways regulated by CC2D1A, such as Akt, CREB and NF-κB, are important for learning and memory (Bourtchuladze et al., 1994; Lai et al., 2006; Majumdar et al., 2011; Meffert et al., 2003). Initial findings in *Cc2d1a*-deficient cells showed an imbalance in signaling activation (Al-Tawashi et al., 2012; Manzini et al., 2014) and mild disruptions in endosome size (Drusenheimer et al., 2015), again demonstrating how CC2D1B is not fully able to compensate for CC2D1A. Our results in the 1a/1b-dHET also imply that there is a balance in CC2D1A and CC2D1B activity, and experiments in Drosophila and mammalian cells suggest that *Cc2d1a* and *lgd* expression and subcellular localization must be finely regulated to control endosomal trafficking and signaling through recruitment to specific signaling complexes (Drusenheimer et al., 2015; Gallagher and Knoblich, 2006; Jaekel and Klein, 2006; Manzini et al., 2014). This could be explained by a critical role for the CC2D1 proteins in the maintenance of signaling homeostasis. Homeostasis is broadly defined as the ability of a cell to return to a set point and maintain equilibrium. Many genes mutated in ASD and ID control homeostatic mechanisms in synaptic transmission, transcription, and signaling (De Rubeis et al., 2014; Pinto et al., 2014), and genomic deletions and duplications may show similar neurodevelopmental phenotypes leading to the hypothesis that pathogenesis of neurodevelopmental disorders is linked to homeostatic imbalance (Ramocki and Zoghbi, 2008). Behavioral impairments in cognitive and social function could then be caused by subtle disruptions in multiple cellular processes limiting the ability of individual neurons and/or neuronal circuits to respond to stimuli, including environmental changes or stressors. In this respect, defining the role of CC2D1A and CC2D1B in homeostatic signaling of multiple pathways disrupted in ASD and ID could be important to dissect whether different signaling pathways contribute to distinct behavioral deficits.

Finally, in light of the cognitive defects in the 1b-KO mice, it may be worthwhile to search for *CC2D1B* mutations in patients with cognitive deficits, and to also consider the possibility of trans-heterozygous cases where *CC2D1A* and *CC2D1B* mutations are both present in heterozygosity. While complete loss of both CC2D1 genes is embryonic lethal, haploinsufficiency of both *CC2D1A* and *CC2D1B* may lead to ID and ASD as *CC2D1A* LOF does. In the Genome Aggregation Database browser, which collects allele frequency data from more than 100,000 individuals in different populations there are 43 likely gene disrupting (stop codon, frameshift or splice site) alleles for *CC2D1A* and 89 for *CC2D1B*. These variants alone or in combination may further contribute to the genetic burden of ID.

## Acknowledgements

The authors are grateful to Tom Maynard and Irene Zohn for advice on mouse line generation and the analysis of the double knockouts, Anthony LaMantia, Judy Liu, Maria Chahrour, Emanuela Santini, and Sally Till for helpful discussion on experimental design. The *Cc2d1a* and *Cc2d1b* KO mouse strains used for this research project were generated by the trans-NIH Knock-Out Mouse Project (KOMP) and obtained from the KOMP Repository (www.komp.org). NIH grants to Velocigene at Regeneron Inc (U01HG004085) and the CSD Consortium (U01HG004080) funded the generation of gene-targeted ES cells for 8500 genes in the KOMP Program and archived and distributed by the KOMP Repository at UC Davis and CHORI (U42RR024244). For more information or to obtain KOMP products go to www.komp.org or.

